# Survey data on European organic multi-species livestock farms

**DOI:** 10.1101/2021.03.24.436791

**Authors:** Defne Ulukan, Lucille Steinmetz, Marie Moerman, Gun Bernes, Mathilde Blanc, Christopher Brock, Marie Destruel, Bertrand Dumont, Elise Lang, Tabea Meischner, Marc Moraine, Bernadette Oehen, David Parsons, Riccardo Primi, Bruno Ronchi, Lisa Schanz, Frédéric Vanwindekens, Patrick Veysset, Christoph Winckler, Guillaume Martin, Marc Benoit

**Author notes:** All three authors contributed equally (Defne Ulukan, Lucille Steinmetz, Marie Moerman).

## Abstract

While there is increasing evidence of the sustainability benefits of diversified systems in the organic cropping sector, this has been much less investigated with organic livestock farming. To fill this knowledge gap, we surveyed a sample of 128 European organic multi-species livestock farms located across seven countries – Austria, Belgium, France, Germany, Italy, Sweden and Switzerland – and covering a large range of livestock species combinations. We recorded 1574 variables as raw data out of which we calculated 107 indicators describing farm structure, management and several sustainability dimensions: resource use efficiency and conservation, animal, land and work productivities, animal and human welfare. After technical validation of the data, we withdrew 26 farms and the database covers 102 farms. This database is well suited to unveil relationships between various dimensions of organic multi-species livestock farm sustainability and their structure and management. It can help reveal sustainable strategies for organic multi-species livestock farming systems and understand levers or barriers to their development.

## Background and summary

The livestock sector is being highly criticized. First, this sector uses 2 billion hectares of pastures and about 700 million hectares of the arable land used for cropping, which is approximately half of the global agricultural area^1^. Livestock also consumes one third of the worldwide cereal production^1^. Using these cereals for meat, milk and egg production is less efficient than their direct consumption by humans, which signifies strong competition between animal feed and human food availability^2,3^. Second, the dominant model of industrial livestock production has well-established direct and indirect impacts on deforestation, climate change, water pollution, soil acidification and biodiversity^4,5^. There is therefore increasing pressure from governments and citizens to step away from this currently dominant model and make more efficient and sustainable use of natural resources^6,7^.

Agroecology is increasingly promoted as a solution to the multiple sustainability issues of world agriculture^8,9^ including in the livestock sector^10^. It entails moving towards more diversified farming systems^11^, i.e. livestock farming systems including multiple breeds of a given livestock species, multiple animal species and even a diversity of crops and pastures. These diversified systems are expected to promote ecosystem services, allowing reductions of input use, to stabilize production levels and income over time^12,13^, and to strengthen farm resilience^14^. While there is increasing evidence of the environmental and economic benefits of diversified systems in the organic cropping sector^15,16^, this has been much less investigated with organic livestock farming.

Multi-species livestock systems are farms where two or more animal species are raised simultaneously. They have received little attention so far^17^. Nevertheless, co-grazing experiments conducted at fine spatial scales (i.e. usually at the level of a field) and over relatively short time horizons (a few weeks) have revealed promising as co-grazing proved to be efficient in natural resource use, while reducing a number of environmental impacts and providing opportunities for animal health management^18–20^. More comprehensive assessments considering the various dimensions of farm sustainability are therefore needed to confirm these promises and provide management opportunities at the farm level. Threshold effects may indeed occur when upscaling experimental outcomes obtained at the field level onto commercial farms.

Motivated by the aforementioned, a survey was conducted in seven European countries between October 2018 and July 2019 that recorded data across 128 multi-species livestock farms. The survey was comprehensive and aimed at gathering data regarding farm structure (farm area, herd size, total number of workers, off-farm activity, etc.), land use (crop and pasture types and areas; management i.e. fertilization, etc.; productivity), livestock management (types of livestock; management i.e. reproduction, diet, housing, health, etc.; productivity), input management (types of products purchased, amounts, etc.), by-product management (types of by-products available, transfers of by-products among farm enterprises, etc.), sales management (on-farm processing, types of product sold, direct selling, etc.), economics (income, satisfaction regarding income) and work conditions (work organization, satisfaction regarding the workload, etc.). Qualitative data on strengths and weaknesses, opportunities and threats perceived by farmers were also collected. The overall database consists of the raw data (1574 variables) and 107 indicators calculated using these variables and reflecting farm structure, management and sustainability of 102 farms. After technical validation, we had to withdraw 26 farms that displayed inconsistent data.

The raw data and the indicators can be used to investigate the relations between farm structure, management and various dimensions of farm sustainability (resource use efficiency, resource conservation, productivity, human welfare, animal welfare) on European organic multi-species livestock farms. It can also serve as a basis to understand the levers and barriers to the development of organic multi-species livestock farming.

## Methods

### Geographic coverage and sampling strategy

The countries in which survey data were collected are represented in Table 1. There are no official statistics available in Europe on multi-species livestock farms, as these are merged with other types of mixed farms in the FADN (2020) database^21^. Thus, it is impossible to know exactly how many multi-species livestock farms exist across Europe. As a result, we did not seek for representativeness of the farm sample but rather tried to explore a diversity of multi-species livestock farms.

**Table 1:**
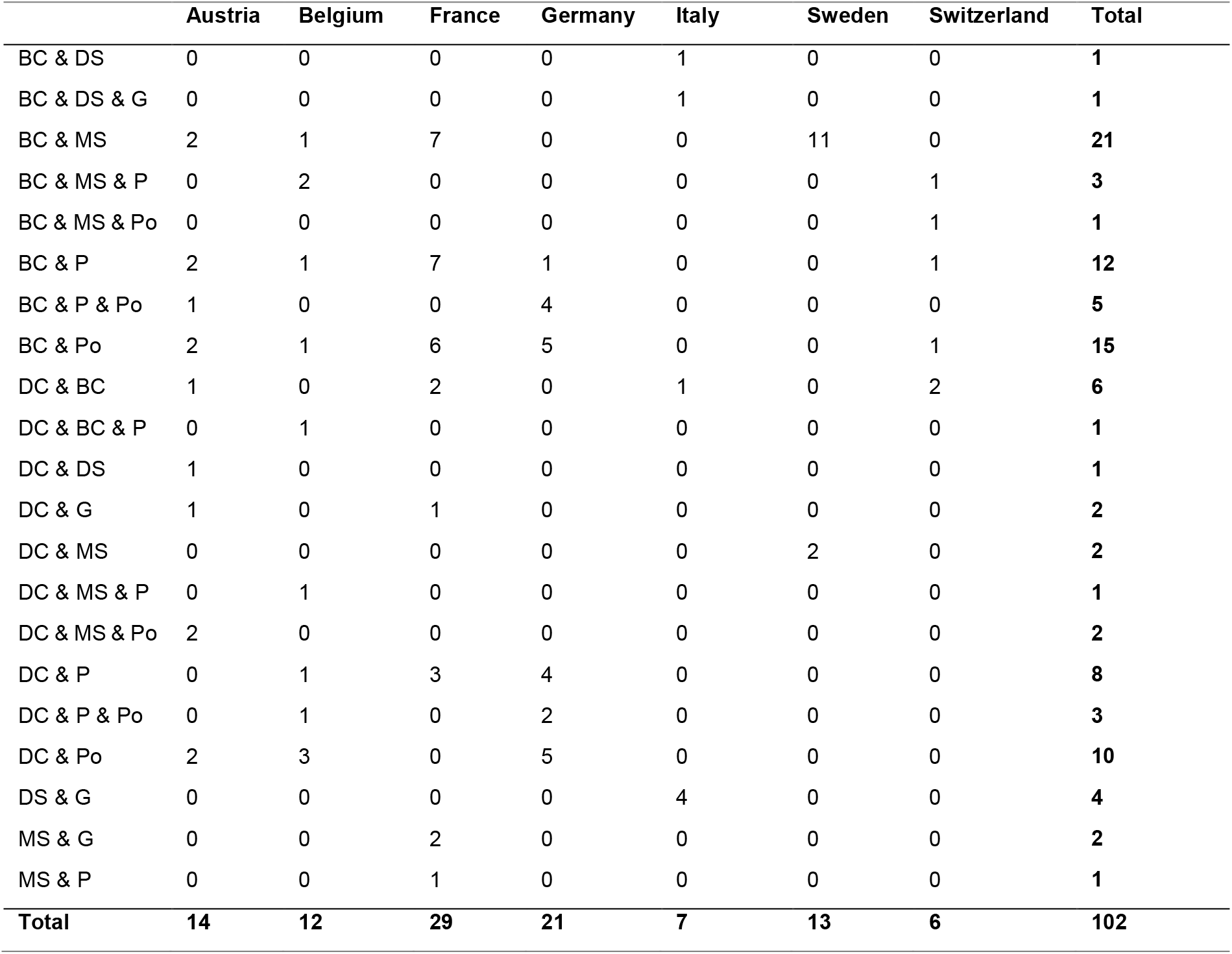
Livestock combinations surveyed per country. BC = beef cattle, DC = dairy cattle, DG= dairy goat, DS = dairy sheep, H = horse, MS = meat sheep, P = Pigs, Po = poultry.

The total number of farms in the final database is 102 (against 128 farms surveyed, more details are provided in the Technical Validation section). The data are all based on individual farm surveys. The farms were chosen from local organic farmer directories, suggestions from local experts or national organic institutions and through snowballing with some farmers suggesting other colleagues. We further applied the following criteria: (i) fully certified organic farms to avoid mixes of free-ranging and confined livestock production activities that do not allow for physical interactions among livestock species; (ii) at least 0.5 worker equivalent unit in the farm to ensure a minimum farm size; (iii) main livestock species limited to cattle, sheep, goat, pig and poultry, with horses and donkeys as possible third species. Additionally, farms with distinct beef cattle and dairy cattle herds were included in the database. This choice was justified in the case of farms conducting those herds as two separate livestock enterprises, thus adding to the diversification of the farm.

### Questionnaire

Through a series of meetings, we developed an ad hoc questionnaire organized into 8 excel sheets addressing farm structure, livestock, pastures, crops, sales, inputs and by-products, economics, and work on the farm. To standardize the survey protocol and the data gathered, most questions called for quantitative, binary or categorical data. Only a few questions and comment boxes were open-ended and allowed farmers to develop their answers. The questionnaire was designed in English and translated into national languages. A pre-test was conducted to test whether the survey guide was understandable for farmers, whether the survey duration was acceptable and whether there were errors that needed to be rectified in the guide before its deployment. Surveys were conducted by researchers and local consultants using this survey guide, in presence of the farm owner(s) or manager. Data covered a full production year and had to be collected for a typical or “average” year, i.e. data smoothed across the years, to clean it from any major climatic, sanitary or economic hazard. Data were first collected in the national languages and then translated in English.

The majority of questions in the survey aimed at allowing the calculation of a series of sustainability indicators. Through these indicators, the following sustainability dimensions were considered:

- Resource use efficiency through output to input ratios enabling verification that farms make efficient use of their inputs to limit environmental impacts;
- Resource conservation through various indicators reflecting the capacity of farms to preserve the natural resources: soils, water, air and biodiversity;
- Self-reliance through technical and economic ratios reflecting the dependence of the farm on its environment;
- Productivity as the production of mega-joules and protein per unit of land, livestock and worker allowing measurement of the contribution of farms to food production;
- Human welfare through farmer satisfaction regarding income, reflecting the capacity of farms to generate decent living conditions for farmers, and farmer satisfaction at work;
- Animal welfare which was assessed through indicators such as livestock mortality.

Sustainability indicators were analyzed in relation to farm structure and farm management. Farm structure accounted for the types of production present on the farms, the natural and human resources available, the climatic and topographical constraints to farming and the diversification activities beyond farming (agritourism, energy production, etc.). Farm management accounted for the management of cropland, pastures, livestock and their interactions via crop rotations, grazing practices, livestock grain or silage feeding, exchanges of by-products, etc. Indicators are also included to represent on-farm processing and sales management (range of products, amounts sold, prices, sale channels, etc.). Work management on the farm was the last component, with indicators illustrating workers’ skills and the distribution of work on the farm.

In accordance with INRAE-Cirad-Ifremer-IRD joint Ethics Committee’s recommendations, our study procedure followed the guidelines provided by INRAE’s Charter of deontology, scientific integrity and ethics^22^. All participants provided informed oral consent prior to the beginning of the survey, after being notified of the purpose of the survey. They were also informed that they had the possibility of skipping questions. Participants did not belong to particularly vulnerable groups. In accordance with the European General Data Protection Regulation^23^, the data were pseudonymised before processing.

### Data processing and indicator calculation

Data for each farm was filled in the questionnaire and saved as an individual excel file. In total, 128 Excel files were compiled and merged into a single database using an R script extracting the data from each individual Excel file. At this stage, the data was pseudonymised. The initial data cleaning was limited to correcting spelling and translation errors, homogenizing character variables when that did not eliminate additional information (e.g. for purchased feed), deleting extra spaces at the end of character strings, making sure missing values were represented by NAs in the database, identifying and correcting obvious impossible values due to forgotten decimal points, wrong units, etc.. This last cleaning step was done with the help of distribution plots for a large number of raw variables. Scripts created in the R software environment were each dedicated to a specific group of tasks. These tasks included processing the raw data on organic multi-species livestock farms and calculating various farm structure, management and sustainability indicators.

Conversion factors used in indicator calculations, such as protein and energy contents of agricultural outputs and inputs, were extracted from public databases (notably, the FoodData Central of the U.S. Department of Agriculture^24^ or the INRAE-CIRAD-AFZ feed tables^25^) or other online literature. They were gathered in correspondence data frames, which linked specific values from our database to the corresponding conversion factor. For instance, a product sold labeled as “cull cows” corresponds to the conversion factor label “cow meat” in the protein and energy tables. This process enabled dealing with a high diversity of labels without losing detailed information in the database itself by creating separate data frames for conversion factors. The final outputs of different indicator calculation scripts were saved in csv-files.

### Data records

The compiled database can be found at: https://doi.org/10.15454/AKEO5G, under the file name “mixenable_data_safe.tab”. Sensitive data, i.e. qualitative data which could be used to identify the farms, were not included in the file to protect farmers’ privacy. All 1574 variables of the survey data are described in the file “Survey data codes.tab”. Variables included in the “Indicator_file.tab” are described in the file “Indicator codes.tab”. Across the entire database, missing values were identified with NA. The “empty_survey_guide.pdf” and the “Guideline to survey Excel file.pdf” are included to provide an understanding of the way the data were gathered. The “empty_survey_guide.pdf” represents the Excel survey file used to record individual farm data and can be read using the zooming in function. R scripts facilitating manipulation of the database and calculation of indicators are also provided (see Code availability).

### Technical validation

Before proceeding to analyses, the data underwent a data quality assessment to ensure its reliability, besides the initial check of impossible values in the raw data. This quality assessment was highly critical for many reasons. First, some variables (technical and economic) were very difficult to collect in a reliable and harmonized way, which required an important phase of consistency checks and adjustments. Second, a minority of farmers provided accounting documents for precisely quantifying inputs and outputs. This was due to the low cultural acceptability of sharing accounting documents in several of the surveyed countries. Consequently, economic variables at farm level were missing for many farms (e.g. gross product, subsidies, debts, etc.) and the economic section of the survey was removed from the final database. Farmer estimation of annual income per associate and farmer satisfaction regarding income were the only variables kept from this section. They were included in human welfare indicators. Lastly, it was also difficult to precisely estimate the stocks of inputs and products at the beginning and end of the production year though these data are essential to calculate reliable technical and economic indicators over a production year^26^.

In the first step, product sales were therefore checked against the resources listed on the farm, i.e. the number of animals in each category, particularly breeding animals for ruminants, and the area dedicated to each crop. The coherence of product sales and animal numbers or crop areas was essential as many indicators were calculated based on these data. As surveys were based on a typical or “average” year, i.e. data smoothed across the years, it was particularly difficult to account for the strong variability of stocks and products sold from one year to another. This was especially true for farms with ruminant meat production as well as for farms with grain production. For instance, a farmer with a suckler cattle herd could keep all 50 calves born in year n for fattening and sale in year n+1. In case all previously born calves were sold during year n-1, there would be no sales during year n. In that case, using data from year n would lead to a dramatic under-estimation of productivity for that farm. As we did not have access to accounting documents for most farms, we opted for correcting sales by stating that the number of animals sold during a year is close to, or slightly lower than, the number of reproductive animals. In cases where there were large discrepancies between estimated and actual sales due to important stock fluctuations, farmers were phone-called to get more information on stock variations and we removed or added animal sales accordingly. Self-consumption was not specifically asked for in the survey, whereas it could have a significant impact on small farm performances and lead to the underestimation of their productivity. It was not possible for us to add an evaluation of self-consumed products but the previously explained consistency adjustments could have helped in reducing this issue. Similarly, the coherence between grain products sold and the area, yield and amounts allocated to sales or on-farm consumption by animals of said grain crop was also checked.

In a second step, to identify inconsistent values, feed purchases and on-farm feed production were checked by comparing them to the theoretical needs of each animal enterprise, considering a large range of variation (lower and upper levels), with the expertise of researchers involved in the project. In order to identify farms where feed consumption was inconsistent, the amount of meat/milk/eggs produced per LU was plotted against the concentrate consumption (purchased + produced) per LU for that enterprise. When on-farm production could not be allocated to fill the gap in concentrate consumption, a purchase of inputs was added in order to obtain the minimum concentrate consumption needed to achieve observed production levels. This was the case when no consumption of feed concentrates occurred, e.g. in milk production or for monogastric animals. Each proposal of adjustment was validated by the farm surveyor.

In a third step, meaningful intermediate output variables were plotted to identify outliers and validate the corrections made in earlier steps. This included plotting for example the total amount of proteins from animal products against the number of LU on the farm (Fig. 2a) and plotting the farm animal productivity (kg protein per total farm LU) against the maximum possible protein production per LU (Fig. 2b). This maximum value capped production to what experts estimated to be physically achievable and allowed the identification of unrealistic productivity values. Caps, i.e. maximum possible values, were based on high animal productivity levels observed in the European context: 10000 kg milk per year per cow, 1400 g daily weight gain for beef cattle, 2 lambs produced (each 50 kg live weight) per year per ewe, 3.1 kg live weight per broiler, 150 kg live weight per pig and 290 eggs per year per hen. These graphs showed that all productivity values were within a reasonable realm for farms kept in the final database.

**Figure 2.**
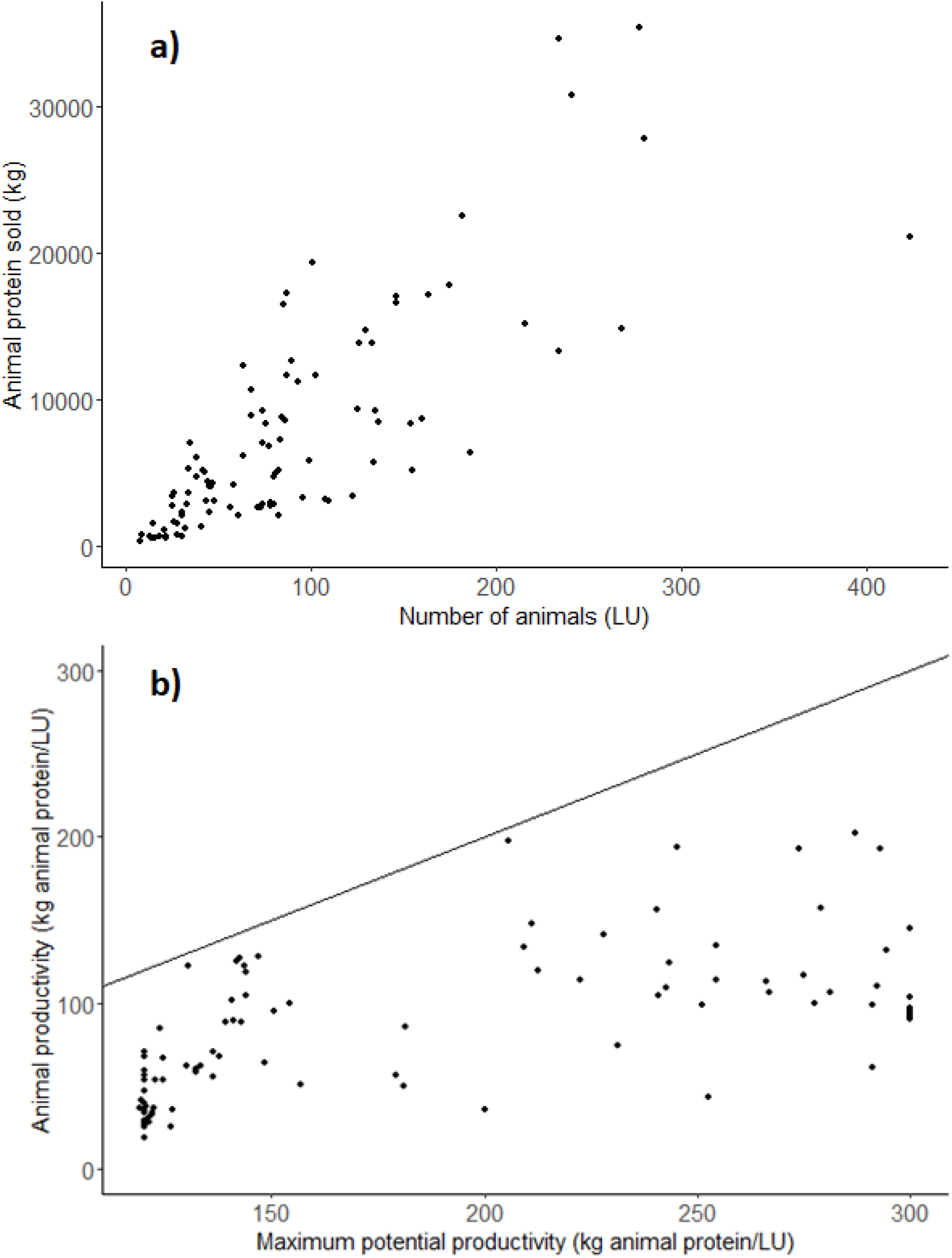
Graphs used for data validation: a) Animal protein sold according to the number of LU at farm level; and b) Animal productivity (in kg of animal protein per LU) depending on the maximum potential productivity (in kg of animal protein per LU). The black line represents the calculated maximum productivity which is achievable for that number of LU.

### Usage notes

The overall strategy that guided the development of this database was to collect farm data offering a first overview of organic multi-species livestock farms at the European level. As explained in the Methods section, this farm sample cannot be considered as representative of the population of European organic multi-species livestock farms which remains unknown. Instead, it covers a diversity of farming contexts (regarding climate, soil, market and regulatory conditions), farms (especially in terms of size and livestock species combination raised) and farming practices (especially regarding those practices determining the level of interactions among livestock species e.g. co-grazing, sequential grazing or grazing on separate pastures). This diversity can eventually be simplified using structural and preferably functional farm typologies^27^.

Against the limited spread of organic multi-species livestock farms, a key issue is to determine the livestock species combinations and management practices (e.g. appropriate stocking rate) especially the level of integration among farm enterprises (e.g. presence/absence of co-grazing, by-product flows among enterprises) required to observe the potential benefits of livestock diversity and avoid undesirable effects (e.g. competition among livestock species at grazing). The database presented in this article is well-suited to such research. It allows identifying farm performance patterns across one to several sustainability dimensions and relating these patterns to explanatory variables including farm structures and farmers’ management practices.

Development of this database was very ambitious because it addressed multiple dimensions of structure, management and sustainability in complex farming systems (diversity of crop and livestock enterprises, of livestock species, of sales channels, etc.). This required gathering a large range of very varied and complementary raw data allowing cross-verification and aiming at understanding farming system coherence. As a result, data collection across several countries turned out to be difficult, leading to an important consolidation and validation phase. Thus, it is important that users consider the survey procedures and the potential biases and limitations mentioned in the technical validation part.

## Acknowledgements

The authors acknowledge the financial support for the MIX-ENABLE project provided by transnational funding bodies, being partners of the H2020 ERA net project, CORE Organic Cofund, and the cofund from the European Commission.

## Author contributions

GM and MBe coordinated the overall work. LSt, MMoe, MD, LSc, GM and MBe designed the survey guide. GB, MBl, CB, BD, DP, RP, PV, CW gave inputs during the design of the survey guide. LSt, MMoe, GB, MBl, MD, EL, TM, MMor, BO, RP, LSc conducted the surveys. DU, MMoe, MD, EL, FV compiled the data. DU, LSt, GM and MBe checked the consistency of the data. DU and GM drafted the manuscript. All co-authors contributed to the finalization of the manuscript.

## Code availability

Scripts and other materials used to calculate indicators can be found at: https://doi.org/10.15454/AKEO5G. Scripts were written under R 4.0.2 through RStudio. They are available in R or Rmd formats. Scripts were named after the theme the indicators they calculate are related to. One script can calculate several indicators around the same theme. External data used in indicator calculations (such as protein content of farm products) were stored in CSV files with a title starting with “References”. The “Initial script” runs all scripts and saves the resulting indicator database.

